# Residual Hair Biomaterial Particulates Improve Dermal Regeneration and Combined with Electrical Stimulation Accelerates Skin Wound Closure

**DOI:** 10.64898/2026.07.04.736482

**Authors:** Dana Saparova, Zara Mahmood, Hope Samuel, Jessica Barayuga, Jash Mody, Nancy Radecker, Roche C. de Guzman

**Affiliations:** Department of Biology, Hofstra University, Hempstead, NY 11549, USA; Bioengineering Program, Department of Engineering, Hofstra University, Hempstead, NY 11549, USA

**Keywords:** wound healing, keratin biomaterial, electrical stimulation, collagen deposition, GAP-43, linear mixed-effects model, AI-assisted image analysis, granulation tissue

## Abstract

**Objective:** To evaluate the effect of residual hair (RH) biomaterial particulates, biphasic electrical stimulation (ES), and their combination (RHES) on the kinetics and quality of skin wound healing.

**Method:** Eighteen adult albino mice received bilateral, splinted 10-mm full-thickness dorsal excisional wounds and were randomly assigned to one of three animal groups producing four wound-level treatment conditions: untreated control (-) (n = 12), RH (n = 12), ES (n = 6), and combined RHES (n = 6 wounds). Daily wound images were segmented using an AI-assisted workflow: a U-Net (ResNet34 encoder, ImageNet-pretrained, trained on a parallel single-expert tracing study with held-out validation Dice = 0.906) generated initial boundary predictions, each reviewed and corrected as needed. Wound size measures (perimeter, area, equivalent diameter [D_eq], circularity, aspect ratio) were normalized to the day-0 value of each wound and analyzed by linear mixed-effects regression with mouse identity as a random intercept and mouse body weight as a covariate. On day 7, wounds were excised, fixed, processed for histology, and analyzed by Masson’s trichrome (collagen content in granulation tissue) and GAP-43 immunohistochemistry (a marker of regenerative cellular activity).

**Results:** All three treatments significantly accelerated wound closure compared to (-) (Day × Treatment interaction χ^2^(3) = 36.4, ***p < 0.0001). The closure-rate advantages on the log-D_eq scale were ES - 0.047/day (***p < 0.0001), RHES -0.029/day (***p = 0.0005), and RH -0.022/day (**p = 0.0015). By day 7, mean D_eq had decreased to 0.58 of the day-0 value in ES, 0.69 in RHES, 0.73 in RH, and 0.79 in (-). Tissue analyses revealed treatment-specific differences in healing quality: RH and RHES wounds contained 6.1× and 8.5× more collagen in granulation tissue than (-) (both **p = 0.002 vs (-); both **p = 0.009 vs ES), and showed approximately 16× and 27× greater mean GAP-43 expression than (-), respectively; the RHES increase remained significant after Bonferroni correction (adjusted *p = 0.042), whereas the RH increase did not (adjusted p = 0.058). ES alone did not significantly increase either collagen content or GAP-43 expression. Wound shape was more circular and more stable across days in RH-containing groups. Mouse body weight did not predict closure, whereas image-derived dryness, eschar coverage, and wound contraction were significant negative predictors of measured wound size.

**Conclusion:** ES, RH, and RHES each significantly improve wound closure kinetics. The improvement appears mechanistically distinct: ES principally accelerates closure rate, while RH principally enhances tissue-level regenerative markers (collagen deposition and GAP-43 expression). RHES combines both advantages.

**Declaration of interest:** Dr. de Guzman is the inventor on a pending patent covering residual hair biomaterial methods and use and a provisional patent on powderized version, both owned by Hofstra University, and is the founder of a company developing wound care products based on the technology. The study was conducted independently. Data are available upon reasonable request.

## 1. INTRODUCTION

Cutaneous wound healing is a coordinated multiphase process whose failure to progress is a major and growing burden on healthcare systems worldwide.^1^ Acute and chronic wounds frequently require interventions beyond standard wound care to reach timely closure, and the search for accessible, low-cost adjuncts remains an active area of investigation. Two strategies of independent biological rationale are biomaterial dressings derived from keratin^2,3^ and exogenous electrical stimulation (ES)^4,5^, each with mechanistic support but limited head-to-head comparison in a model designed to suppress contraction and reflect true tissue repair.

Residual hair (RH) biomaterial consists of organized, keratin-rich fibers and macrofibrils prepared from the insoluble fraction of processed human hair waste.^6,7^ Keratin biomaterials present structural and biochemical cues associated with epithelial migration, exudate management, and granulation tissue formation, and have shown clinical benefit on refractory wounds.^8^ Biphasic micro-current ES reinforces the endogenous bioelectric fields generated immediately after cutaneous injury and has been associated with directional keratinocyte migration, reduced inflammation, and improved tissue perfusion.^9–11^

In a preceding pilot study using the same bilateral splinted murine excisional model with a pure RH extract of short strands and a sub-nominal ES current, our group observed only limited closure-rate advantages and no tissue-level benefit. Two adjustments were therefore implemented for the present study: an excipient-tuned, fine-particulate RH formulation intended to improve moisture balance, surface area, and adherence, and verified delivery of the targeted ES current. We additionally implemented an AI-assisted boundary-tracing workflow and quantitative analyses of histology and immunohistochemistry to capture treatment-specific effects on healing quality.

This paper reports a 7-day evaluation of the kinetics, shape, and tissue-level outcomes of wound healing across four treatment conditions in 18 mice. We hypothesized that each treatment, alone or combined, would improve wound healing over untreated controls but through partially distinct mechanisms detectable when both kinetic imaging and endpoint histology are assessed in the same animals.

## 2. METHODS

### 2.1 Residual hair biomaterial particle preparation and characterization

Human hair clippings were collected from anonymized donations from local Long Island, NY salons. Natural dark/black strands ≥1 inch were retained while naturally-blonde and bleached strands were excluded. Hair was cleaned in 0.2% sodium dodecyl sulfate, rinsed, and air-dried. Melanin was oxidized by 24-h treatment of 1 part hair with 2 parts 40-volume developer and 1 part lightener, followed by extensive water washing. Matrix proteins (keratin associated proteins) were depleted with two rounds of tris/ethanol/thioglycolic acid extraction^6^ and phosphate buffered saline (PBS) neutralization to concentrate organized macrofibrils and keratin intermediate filaments in the insoluble residual fraction. Cuticles were weakened by three 10-min cycles of boiling in 88% formic acid (in a fume hood), followed by PBS soaking, water washing, and oven drying at 80 °C for 15-20 h. The residual hair (RH) was cryo-particleized by liquid-N_2_ mortar-and-pestle grinding (10-15 min). Another batch of RH was powderized in a mechanical grinder for 3 min, repeated thrice. Samples were visualized in scanning electron microscopy (SEM, sputter-coated with gold, Q250 FEI at 30 kV). Size profiles were evaluated using ImageJ (NIH). This second batch (with finer particles) was formulated with excipients to support hydration into a paste-like consistency, for subsequent animal wound treatment application.

### 2.2 Electrical stimulation device

The biphasic ES device (Biothm, Prowell Technology) was configured to a higher amplitude than that used in our preceding pilot study, with target current ∼ 88 μA, phase width 350 μs, and frequency 20 Hz. Output was verified by multimeter measurement of voltage across the wound-gel pad electrode interface (12 × 17 × 1 mm pads) before each session. MATLAB (MathWorks) was employed to simulate the biphasic electric current output as a function of time to demonstrate the phase width (PW) signals at the beginning of the period and to show a pulse train consisting of multiple periods.

### 2.3 Animal study

The Hofstra University Institutional Animal Care and Use Committee approved this animal study. Eighteen adult albino mice (8-12 weeks, ≥ 25 g; mouse IDs: a–r) were randomly assigned to one of three animal groups ((-)/ES, RH/RHES, and (-)/RH), producing 36 bilateral wounds across four wound-level treatment conditions: untreated (-) n = 12, RH n = 12, ES n = 6, and combined RHES n = 6. Three male (M) and three female (F) mice were assigned per animal group. Wound treatment side (L = left vs R = right) followed a fixed convention per animal group as shown in **Table 1**.

**Table 1.** Treatment assignment by animal group.

| Animal group | (Sex: IDs) | Treatment |  |
| --- | --- | --- | --- |
|  |  | Left | Right |
| (-)/ES | (<u>F</u>: a, b, c & <u>M</u>: d, e, f) | (-) | ES |
| RH/RHES | (<u>F</u>: g, h, i & <u>M</u>: j, k, l) | RH | RHES |
| (-)/RH | (<u>F</u>: m, n, o & <u>M</u>: p, q, r) | (-) | RH |

Mice were anesthetized with isoflurane (0.6-1% in filtered air, SomnoSuite, Kent Scientific). Dorsal hair was shaved and depilated. Two 10-mm circular full-thickness wounds were created with a sterile biopsy punch followed by complete excision with iris scissors. Each wound was stabilized with a 0.8 mm thick annular silicone splint^12^ (10 mm ID and 15 mm OD) affixed by four cardinal-direction drops of tissue adhesive (LiquiVet Rapid, Oasis Medical). For RH-receiving wounds, RH formulation was applied to the wound bed (approximately 50 μL). RH was reapplied daily as needed. Wounds were covered with a polypropylene-polyester occlusive dressing (Opsite Flexifix, Smith & Nephew). Animals received ketoprofen (5 mg/kg) post-surgery and for 2 additional days.

Under daily anesthesia, dressings and splints were carefully removed (not reinjuring the wound) for treatment and imaging. ES was applied once daily from day 1 to day 6 at 20 Hz biphasic, 350 μs phase width, target ∼ 88 μA, 5 minutes per session per modified parameters from Sari et al.^13^. Wounds were imaged with a stereomicroscope at 0.63× objective (Stemi 508, Zeiss) at 1920 × 1080 pixel resolution (covering the whole 10-mm wound in the field of view). New splints and occlusive dressings were reapplied after imaging. A total of 288 top view images were collected from 3 animal groups × 6 biological replicates (3 M and 3 F) × 2 wound locations (L and R) × 8 time points (day 0 to day 7). On day 7, animals were euthanized, the wound and surrounding skin excised, and the tissue divided: one half was snap-frozen for future molecular analysis and the other was processed for histology.

### 2.4 AI-assisted wound boundary segmentation

A U-Net convolutional neural network (CNN) with a ResNet34 encoder pretrained on ImageNet (segmentation-models-pytorch 0.5.0) was trained on a parallel single-expert (Dr. de Guzman) dataset of 274 dorsal wound images polygon-traced in ImageJ. Training used 512 × 512 pixel input, combined binary cross-entropy and Dice loss, AdamW optimizer (learning rate 1 × 10^-^L, cosine annealing) for 40 epochs, batch size 8, on a Tesla T4 GPU (Google Colab). Augmentation included horizontal/vertical flips, 90° rotation, affine ± 20°, elastic deformation, and color jitter. Train/validation (80/20) split was stratified by treatment and split by wound (no validation-set image came from a training-set wound).

The trained model was then used to generate the initial boundary predictions for all 288 top-view wound images. Each prediction was visually reviewed by the tracer on per-wound review pages displaying all eight days (day 0 to day 7) side-by-side. Predictions matching the wound margin were ACCEPTED and the auto-derived measurements (area, perimeter, major/minor axis, circularity, D_eq) extracted directly. Predictions requiring adjustment were classified CORRECT (boundary refinement needed) or REJECT (model failed entirely) and re-traced manually in ImageJ using the same convention. All final measurements thus passed expert visual validation.

Per-image observation covariates were scored by the same tracer for each image: TissueGlue (binary, presence of tissue adhesive on the wound), Bleeding (binary, presence of blood), Hair (binary, presence of regrowing hair), Exudate (0-3 ordinal, amount of exudate), EscharCoverage (0-3 ordinal, degree of eschar), Dryness (0-2 ordinal, wound bed drying), Contraction (binary, evidence of wound contraction), Distorted (binary, indicating visible distortion of the wound by loose skin during imaging), and Quality (1-3 ordinal, image clarity and quality).

### 2.5 Quantification of wound size and shape

From the boundary polygon for each image, the following measurements were measured: area (A), perimeter (P), major and minor axes of a fitted ellipse, Feret’s diameter, equivalent diameter: D_eq = 2√(A/π), circularity = 4πA/P^2^, and aspect ratio = major/minor. The D_eq is used as the primary outcome because it is derived from area (and is therefore less sensitive to local perimeter roughness from eschar or biomaterial) yet expressed in linear units (and therefore less sensitive to imaging-angle foreshortening than area). Perimeter and area ratios are reported as concordant secondary outcomes. Each measurement was normalized to its own day-0 value, so that every wound begins at a ratio of 1.0 and subsequent change reflects closure relative to the initial wound.

### 2.6 Histology and immunohistochemistry

From each of the 36 wounds, six wounds per treatment condition were systematically selected for histology (three M, three F per group, randomly assigned at study start). Each wound contributed two adjacent sections: one stained with Masson’s trichrome (MT) and one for immunohistochemistry (IHC). All histological and immunohistochemical imaging was performed using identical microscope settings across experimental groups to ensure comparability of signal intensity and tissue morphology.

Following tissue harvest, samples were fixed in 10% neutral buffered formalin and processed through graded ethanol dehydration, xylene clearing, and paraffin embedding. Embedded tissues were sectioned at approximately 5 µm thickness using a rotary microtome and mounted onto glass microscope slides. Sections were stained using Masson’s trichrome stain according to standard histological protocols.^6^ MT-stained sections were imaged at 4× under transmission microscopy (Zeiss) and merged across the wound site and adjacent intact skin. Collagen content in granulation tissue was quantified in ImageJ as the fraction of pixels above a blue-channel threshold within a manually-traced granulation tissue ROI (collagen / granulation tissue area). A few sections were unreadable or damaged during processing and were excluded (final n per group: (-) n = 6, ES n = 5, RH n = 6, and RHES n = 6).

For assessment of cutaneous nerve regeneration, IHC was performed using growth-associated protein 43 (GAP-43), a marker associated with axonal sprouting and regenerating nerve fibers.^14^ Tissue sections underwent deparaffinization, rehydration, antigen retrieval, and blocking of non-specific binding prior to incubation with the primary anti-GAP-43 antibody (Rabbit GAP43 Polyclonal Antibody, bs-0154R, Bioss). Following incubation with the appropriate fluorescent secondary antibody (Goat Anti-Rabbit IgG Antibody (H+L), AbBy Fluor^®^ 488 Conjugated, bs-0295G-BF488, Bioss), sections were counterstained and mounted with antifade medium with DAPI nuclear stain. Samples were imaged using a laser scanning confocal microscope (FV3000, Olympus). Relative GAP-43 expression was computed as the integrated 488-channel signal divided by the tissue area within an ROI placed in the dermal granulation tissue. Final n per group after exclusion of unreadable/damaged sections: (-) n = 4, ES n = 6, RH n = 5, and RHES n = 6.

### 2.7 Statistical analysis

Wound size (log D_eq ratio) was modeled by linear mixed-effects regression in Python (statsmodels): log(D_eq,day / D_eq,day0) ∼ Day × Treatment + Sex + Side + Weight + covariates + (1 | Mouse) with Treatment categorical (reference = (-)), mouse identity as a random intercept, and mouse body weight included as a centered fixed-effect covariate. The Day × Treatment interaction tests treatment-specific closure rates. Additional per-image covariates entered the model as fixed effects: TissueGlue, Bleeding, Hair, Exudate, EscharCoverage, Dryness, Contraction, Distorted, and Quality. Day 0 was excluded from estimation because its ratio is 1.0 by construction. Days 1-7 were used for the closure-rate analysis. Models were fitted by Restricted (or Residual) Maximum Likelihood (REML), and fixed-effect inference used joint Wald χ^2^ tests per term. Equivalent models fitted on log-perimeter and log-area ratios served as concordant sensitivity analyses. Wound shape (circularity and aspect ratio) and within-wound shape stability (the SD across days) were summarized descriptively and compared between treatment groups by the Kruskal-Wallis test. Day-7 endpoint tissue measures (Masson’s trichrome collagen fraction and GAP-43 expression, n = 4-6 per group) were compared by Kruskal-Wallis followed by pairwise Mann-Whitney U tests versus (-), with Bonferroni adjustment for the three treatment-versus-control comparisons. Significance reported as: *** p < 0.001, ** p < 0.01, * p < 0.05.

## 3. RESULTS

### 3.1 Two versions of RH particulates and their characterization

In the first process using liquid-N_2_ manual grinding, the RH particulates look powdery in appearance with yellow to light brown in coloration (**Fig. 1a**). It accumulates considerable static electricity, similar to the soluble version of human hair extract, kerateine.^15^ Under SEM (**Fig. 1b**), at lower magnification, the particles appear uniform but upon zooming in to higher magnification, the cylindrical residual hair structure was mostly preserved (Fig. 1b). In some high magnification images, the internal cortical structures containing macrofibril and intermediate filament organization of keratins were evident (**Fig. 1b, arrows**). The outermost cuticle appears to have been eroded. The glassy appearance suggests that water is still present in the matrix of the dried RH particles. The length of ground RH did not follow a normal distribution (Fig. 1d). This suggests that the process is still incomplete since the length’s standard deviation needed to be brought to 1/3 of its current value to reach the expected normal distribution. The diameter follows the normal distribution (Fig. 1e). Particle properties: length 122 ± 104 μm and diameter 58 ± 16 μm.

**Figure 1.**
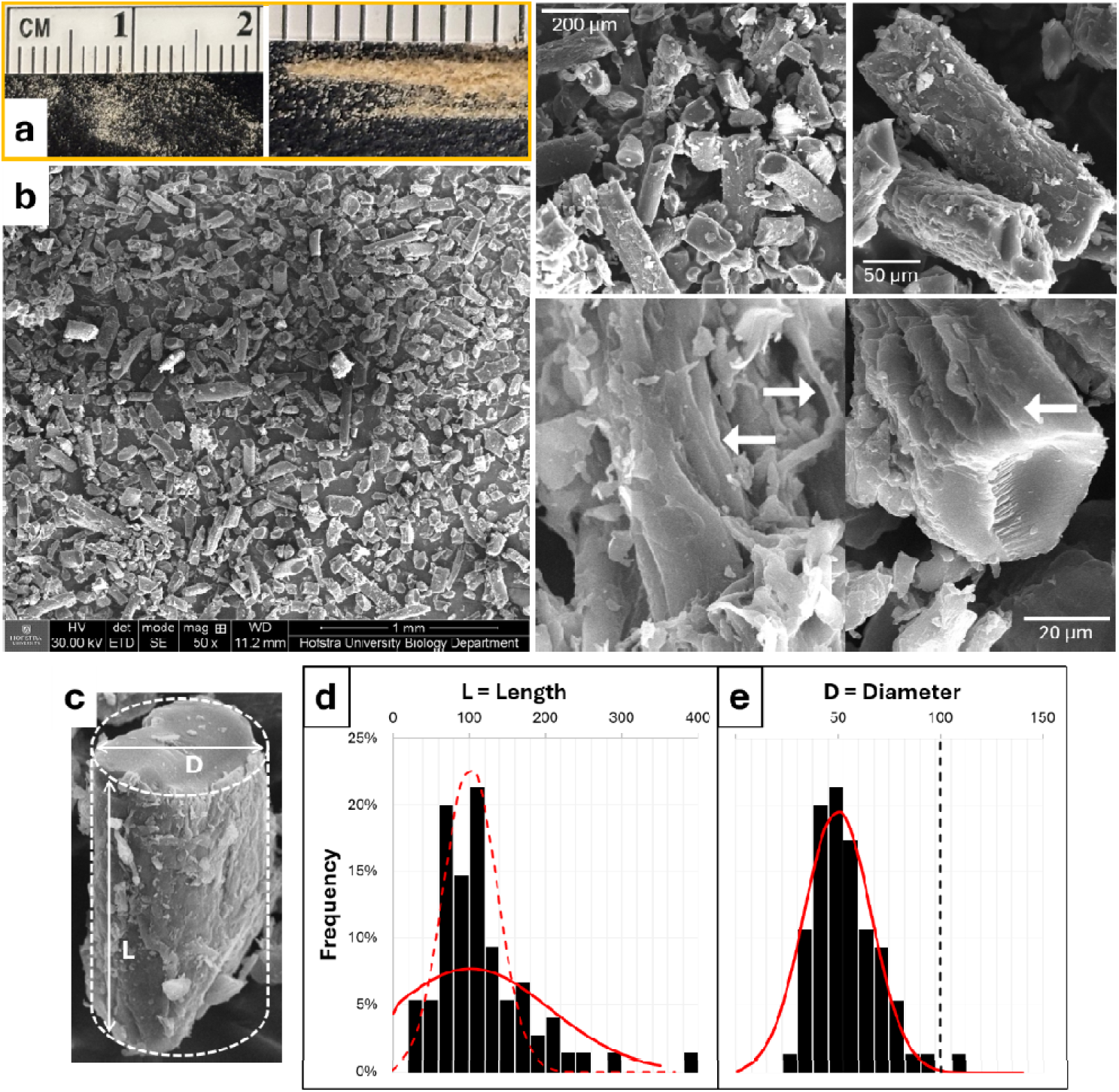
Liquid nitrogen-ground residual hair biomaterials imaged via **a)** macro light, and **b)** SEM at increasing magnification. Arrows show the presence of aligned internal structures, representing macrofibrils and intermediate filaments containing macromolecularly-organized hair keratin proteins in exposed internal cortex of hair particulates. **c)** The particles are assumed to be cylinders with length (L) and diameter (D). **d)** The length distribution does not follow a normal distribution (red dashed), but ideally when the standard deviation is decreased to 1/3 its current value then it will obey the normal distribution (red solid curve). **e)** The diameter histogram (bar) follows normal (red solid curve) with mean at 58 μm, about half of original/untreated hair’s diameter (black dashed).

Using a machine grinder resulted in a bimodal distribution of particles (**Fig. 2a**): the bigger ones with length 84 ± 43 μm (**Fig. 2b**) and diameter 62 ± 30 μm (**Fig. 2c**), and a smaller ones with length 21 ± 9 μm and diameter 14 ± 6 μm. Particle sizes followed a normal distribution. When formulated with excipients, it appears yellowish (**Fig. 2d**) and zooming in microscopically shows white hydrogel structure. The RH formulation is spreadable on the surface of an excised mouse skin wound (**Fig. 2e**).

**Figure 2.**
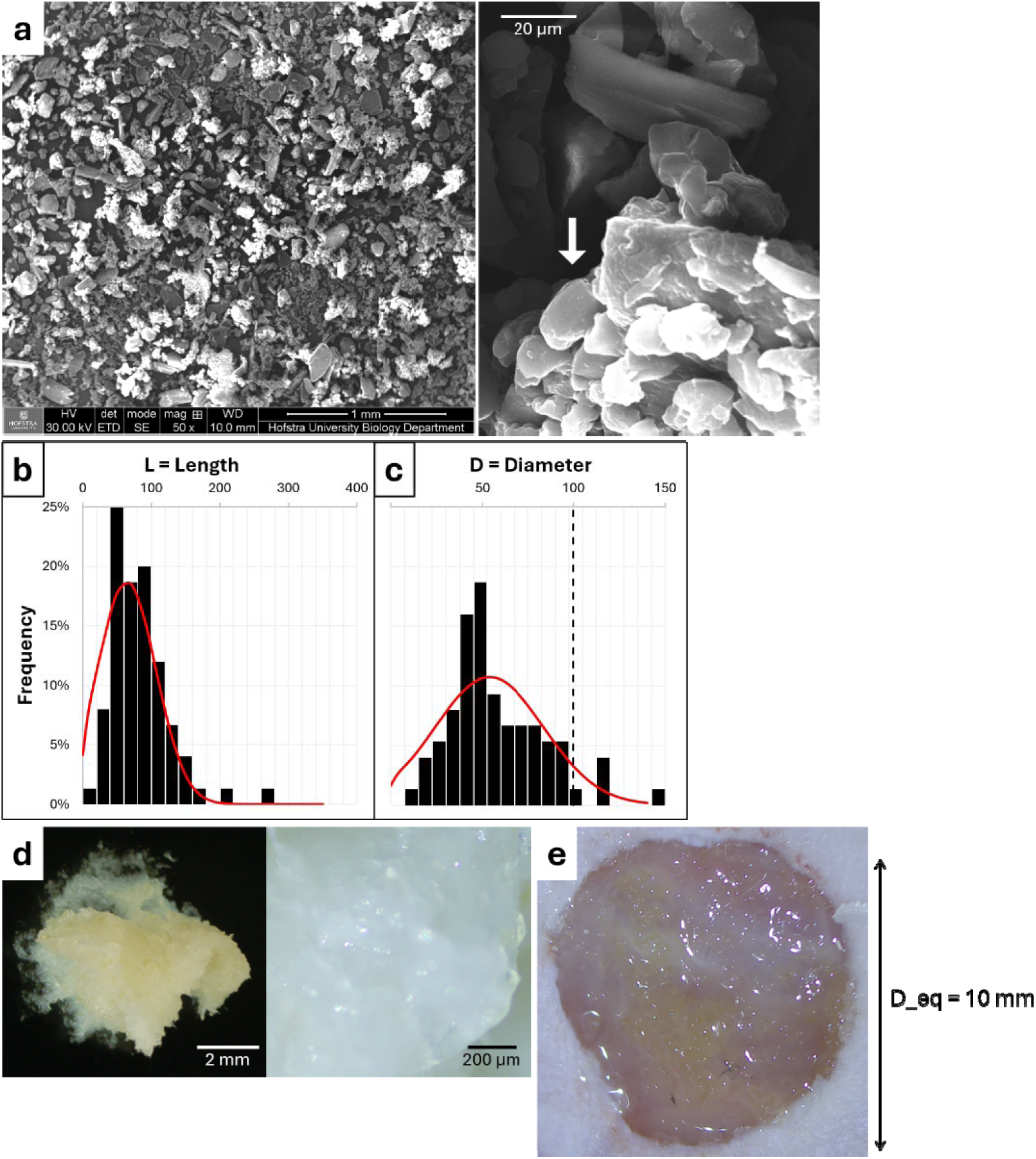
**a)** Machine-ground residual hair biomaterials produce two distinct sizes of particles: the bigger preserving the hair structure with normally-distributed **b)** length and **c)** diameter, and the smaller (arrow) cortical bundles of macrofibrils. **d)** Ground residual hair mixed with excipients under stereomicroscope and **e)** when applied to a representative 10-mm diameter mouse excision skin wound.

### 3.2 Electrical stimulation device testing

The ES device outputted the expected I = electric current amplitude at ∼ 88 μA and f = frequency at ∼ 20 Hz (**Fig. 3a**) when connected directly to the metallic electrodes and gel pads. The PW of 350 μs was not tested due to the very small time scale. Simulation showed (**Fig. 3b**) that the initial dip in current (negative I) is the cathodic followed by an increase to positive I (anodic), each corresponding to PW, and double the PW is the stimulation pulse at 700 μs = 0.7 ms. The remaining period is zero A up to the period of T = 50 ms. The duty cycle was computed to be at 0.7 ms/50 ms = 0.014 = 1.4%. The pulse train containing multiple periods (4 cycles) was simulated over 200 ms (**Fig. 3c**), demonstrating that the ES signals are biphasic spikes at a high frequency of 20 Hz. Physical testing on the live anesthetized mouse with excision wound (**Fig. 3d**) using an ammeter configuration over multiple trials at the target 5-minute interval confirmed the target wound is receiving ∼ 88 μA of electric current.

**Figure 3.**
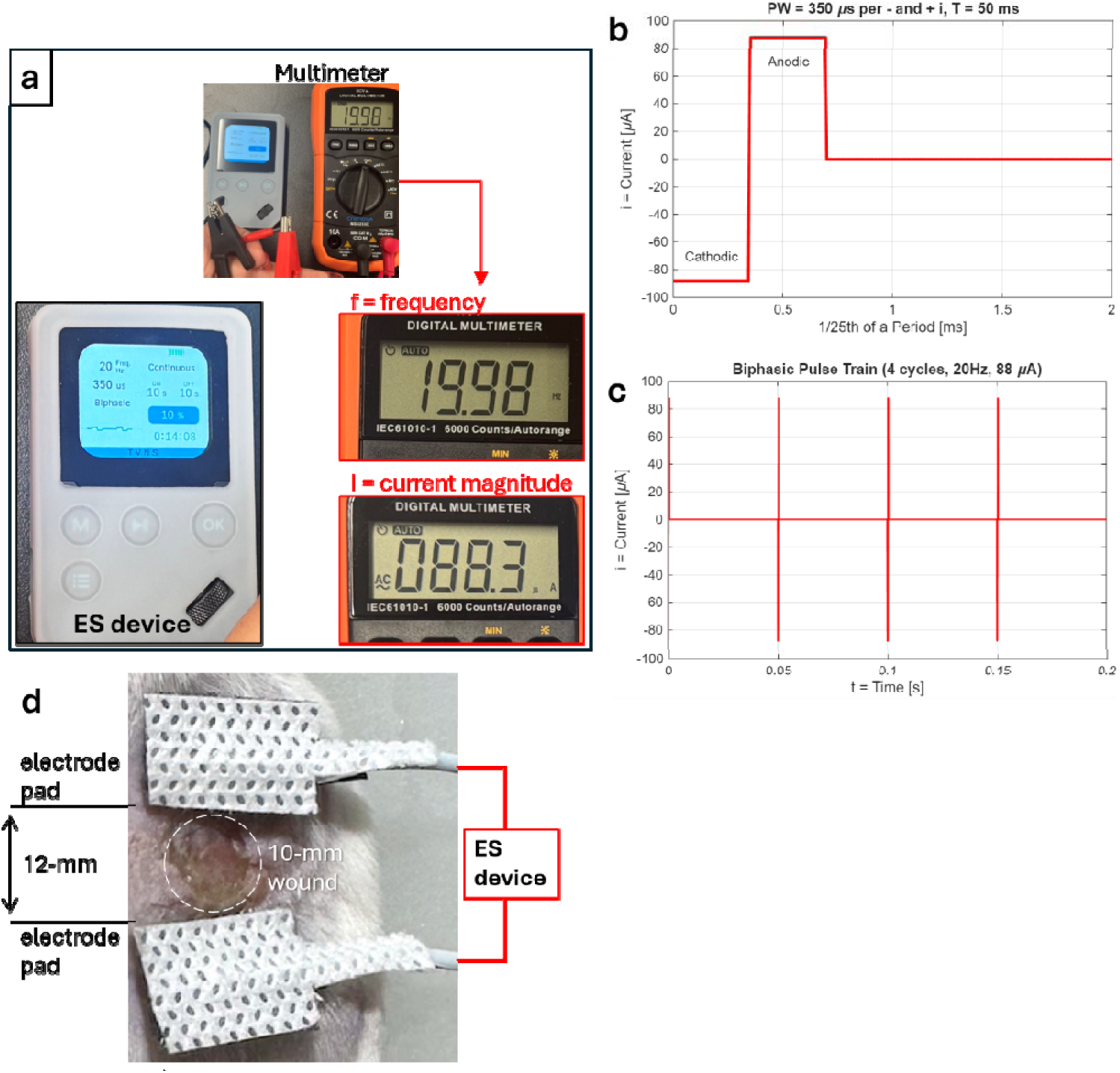
**a)** The ES device is programmed and tested using a multimeter for outputs including frequency and electric current magnitude. MATLAB simulation of the waveforms showing: **b)** a section of the period (1/25th) encompassing the phase width (PW) for biphasic: both the negative (cathodic) and positive (anodic) pulses and **c)** a 4-period pulse train. **d)** Gel pads connected to the electrodes were separated by approximately 12-mm, encompassing the 10-mm diameter wound on the right side of the mouse. ES was applied on anesthetized mouse daily for 5 min.

### 3.3 Machine learning model outcome

The U-Net CNN-ResNet34 training/validation model reached a best held-out validation Dice coefficient of 0.906, with a median wound-area error of 6.1% and a median perimeter error of 10.6% relative to manual tracing, indicating close agreement with expert segmentation.

When employed on wound top view 288 images, 243 (84.4%) were ACCEPTED, 38 (13.2%) CORRECTED, and 7 (2.4%) REJECTED. A representative auto-traced boundary is shown in **Fig. 4**.

**Figure 4.**
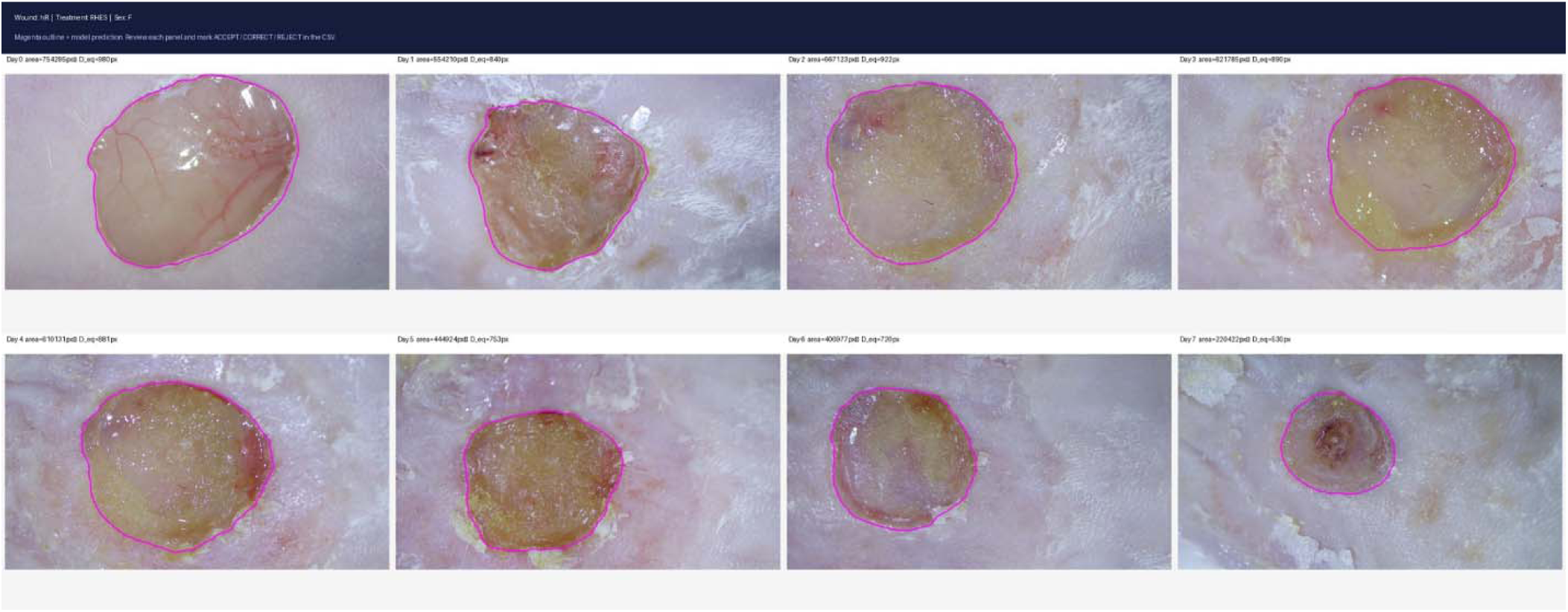
Representative top-view wound auto-tracing output (magenta) from the residual hair (RH) particulate formulation group based on the custom-trained machine learning tool.

### 3.4 Wound closure kinetics

Wound sizes progressively decreased over the one-week span, showing skin wound healing across all treatment groups (including the untreated (-) control) (**Fig. 5**). Using the machine learning auto-traced wound boundaries, equivalent diameter (D_eq) was obtained (along with secondary size metrics: perimeter and area). When normalized to respective day 0s, the quantified values demonstrated wound size decrease for the concordant size metrics (perimeter, area, and D_eq) (**Fig. 6**). By day 7, the mean D_eq had decreased to 0.79 in the untreated (-) group, 0.73 in RH, 0.69 in RHES, and 0.58 in ES: i.e., the ES group showed the greatest closure (42% reduction) while the (-) group the least (21% reduction).

**Figure 5.**
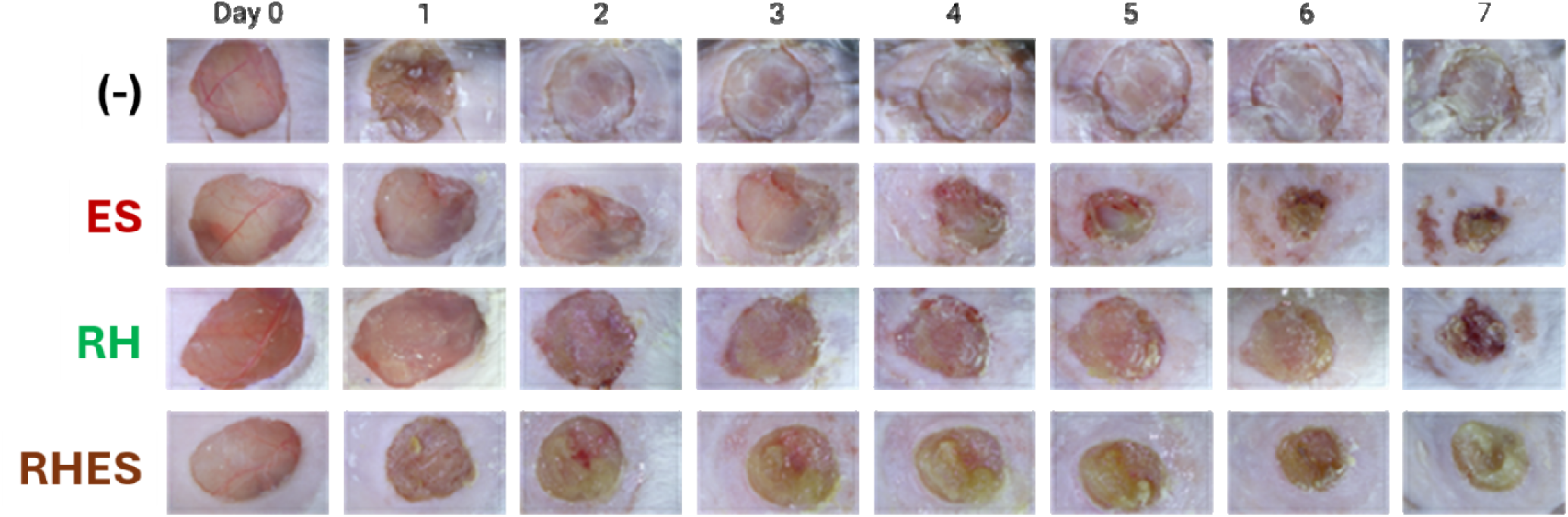
Representative top view wound images over time (days 0 to 7) from the four different treatment groups ((-), ES, RH, and RHES).

**Figure 6.**
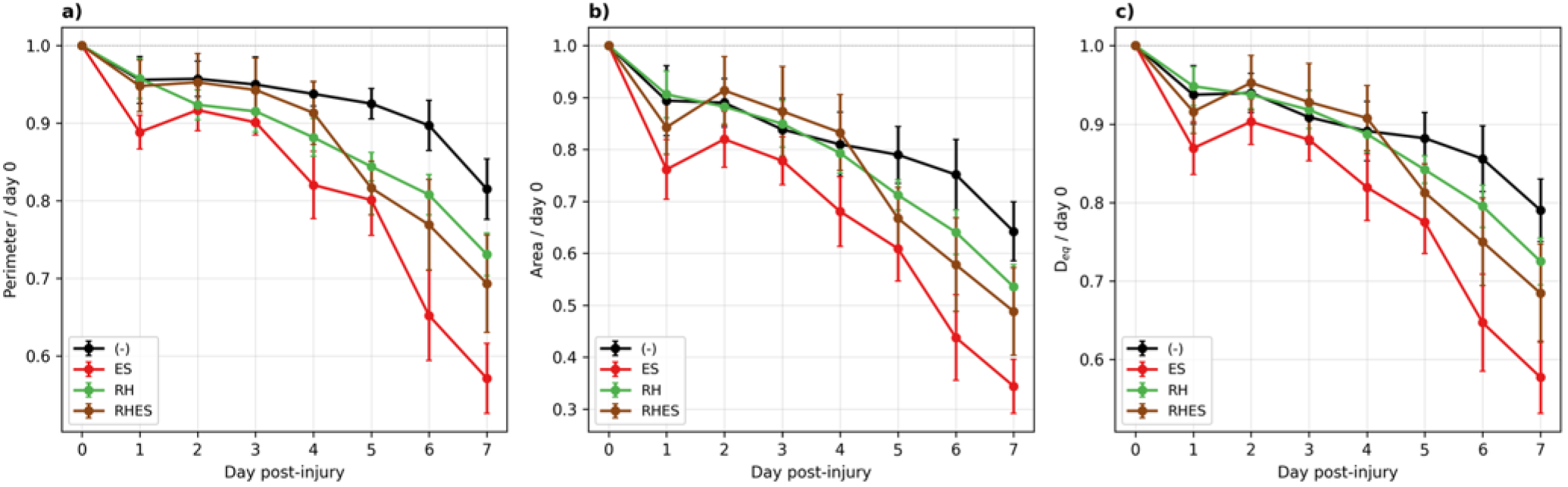
Wound size trajectories by treatment for three concordant size metrics: **a)** perimeter, **b)** area, and **c)** equivalent diameter, each normalized to the day-0 value (mean ± SEM across wounds). All three metrics show the same ordering: ES achieves the greatest closure by day 7, followed by RHES, RH, and (-). Sample sizes per treatment: (-) n = 12, RH n = 12, ES n = 6, and RHES n = 6.

The linear mixed-effects model identified a highly significant Day × Treatment interaction (χ^2^(3) = 36.4, ***p < 0.0001 on log-D_eq), confirming that closure rate differed by treatment. The overall time effect (Day, joint test) was also strongly significant (χ^2^(4) = 149.2, ***p < 0.0001), validating the progressive closure across all groups. Pairwise treatment-versus-(-) coefficients on the log-D_eq scale (**Table 2**) showed ES with the largest closure-rate advantage (β = -0.047/day, ***p < 0.0001), RHES second (β = -0.029/day, ***p = 0.0005), and RH third but still significant (β = -0.022/day, ***p = 0.0015). The interaction estimates were qualitatively concordant on the log-perimeter and log-area scales. Mouse body weight, included as a centered fixed-effect covariate, was not a significant predictor of wound size (main effect p = 0.70) or of closure rate (weight × day interaction p = 0.51), and sex and wound side were likewise non-significant, confirming that closure kinetics were independent of animal size, sex, and anatomical side.

**Table 2.** Linear mixed-effects analysis of log(D_eq ratio), normalized to day 0, for days 1-7. Coefficients on Day × Treatment terms represent the additional change in log-D_eq per day relative to the untreated (-) group. Negative coefficients indicate faster closure than control. Mouse body weight was included as a centered covariate. The remaining covariates (TissueGlue, Bleeding, Hair, Exudate, Contraction, Distorted, Quality) did not reach statistical significance.

| Term | $\chi^2$ / coefficient | df | p-value | Significance |
| --- | --- | --- | --- | --- |
| Day (overall time effect, joint test) | 149.16 | 4 | < 0.0001 | *** |
| Treatment (main effect) | 1.41 | 3 | 0.704 | NS |
| Day × Treatment | 36.36 | 3 | < 0.0001 | *** |
| Day × RH (vs -) | $\beta = -0.022/\text{day}$ | 1 | 0.0015 | ** |
| Day × ES (vs -) | $\beta = -0.047/\text{day}$ | 1 | < 0.0001 | *** |
| Day × RHES (vs -) | $\beta = -0.029/\text{day}$ | 1 | 0.0005 | *** |
| Sex | 2.11 | 1 | 0.146 | NS |
| Side | 0.27 | 1 | 0.603 | NS |
| Dryness covariate | 32.89 | 1 | < 0.0001 | *** |
| EscharCoverage covariate | 7.04 | 1 | 0.008 | ** |
| Mouse weight (joint test: main + weight × day) | 0.71 | 2 | 0.703 | NS |
| Contraction covariate | 4.75 | 1 | 0.029 | * |

Three per-image covariates were significant negative predictors of measured wound size: Dryness (***p < 0.0001), EscharCoverage (***p = 0.008), and Contraction (*p = 0.029). Each reduces the measured wound-size ratio when present, consistent with passive contraction following dressing failure (Dryness), obscuration of the wound margin by overlying crust (EscharCoverage), and visible wound-margin contraction during the early proliferative phase (Contraction). Importantly, including these covariates did not materially change the Day × Treatment estimates, indicating that the treatment effects are robust to image-level artifacts.

### 3.5 Wound shape

Wound circularity (4πA/P^2^) and aspect ratio (major/minor axis) were tracked across the 7-day period (**Fig. 7a-b**). RH-containing wounds maintained slightly higher mean circularity (RH 0.77, RHES 0.78) than untreated controls (0.73), with ES intermediate (0.77). Aspect ratio was correspondingly lower (more circular) in RH (1.24) and RHES (1.28) than in (-) (1.31). More notably, within-wound shape stability, the standard deviation of circularity across all days for each wound, was lowest in the RH-containing groups (median SD 0.036 for RH and 0.027 for RHES) and highest in untreated controls (0.052), with ES intermediate (0.037) (**Fig. 7c-d**). These differences are consistent with a mechanical-stabilization interpretation in which the RH biomaterial layer dampens passive distortion of the wound bed and constrains anisotropic granulation, yielding more uniform wound geometry over time.

**Figure 7.**
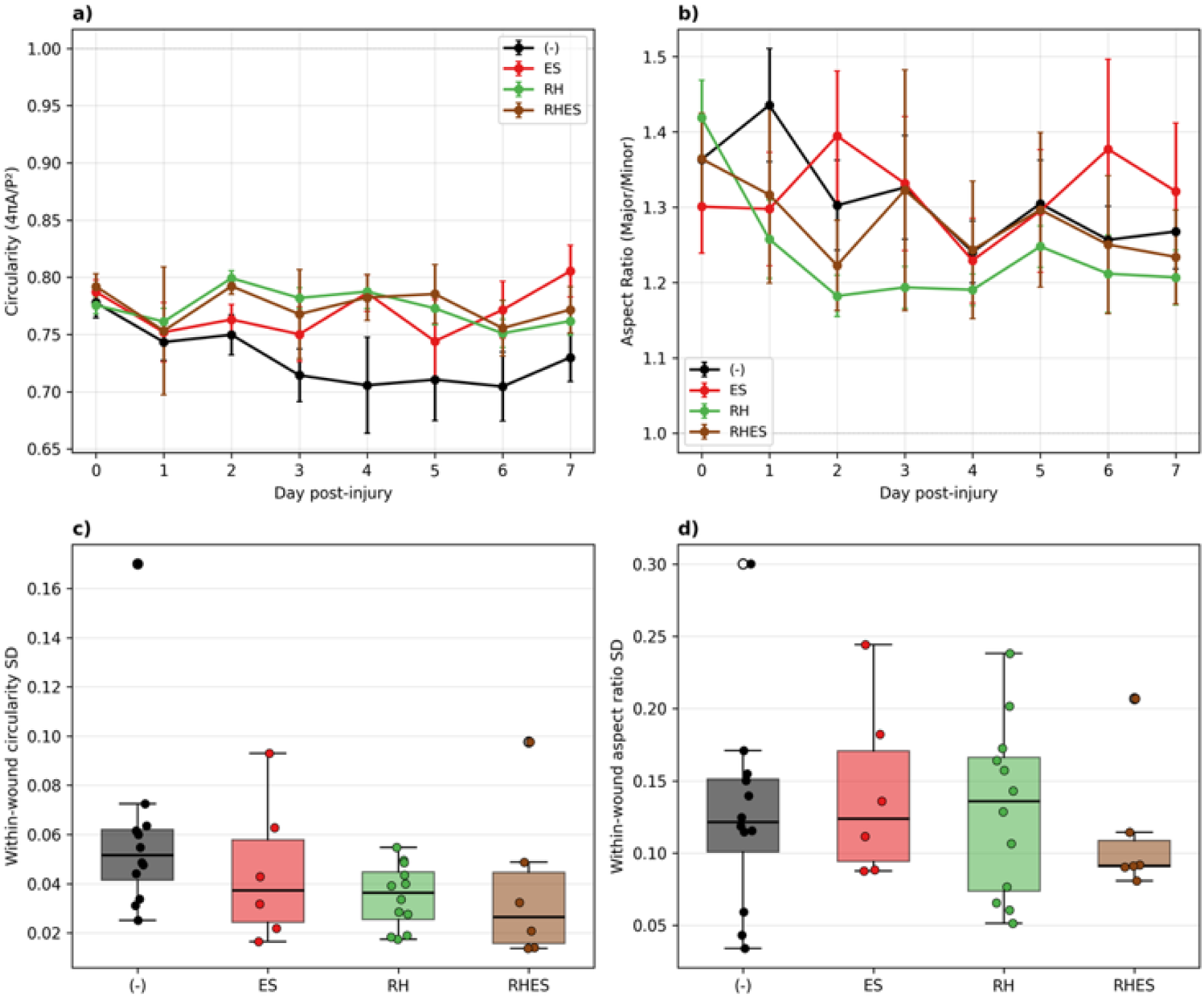
**a)** Wound circularity and **b) aspect ratio over time**, by treatment (mean ± SEM). RH and RHES wounds are consistently more circular and less elongated than untreated controls. Within-wound shape stability, summarized as the standard deviation across the 7-day period for each wound. RH and RHES wounds show the lowest within-wound variability in both **c)** circularity and **d)** aspect ratio, indicating mechanically more stable wound geometry. Box: median and interquartile range; whiskers: 1.5× IQR; points: individual wounds.

### 3.6 Underside blood vessels at day 7 (qualitative observation)

At day 7, the excised wound and surrounding skin were inverted and the underside imaged to assess the dermal and subdermal vasculature supplying the healing wound bed. On qualitative inspection, the wound undersides of the ES- and RHES-treated groups generally displayed denser and more branched networks of fine blood vessels converging on the wound margin than the untreated controls, while RH undersides appeared comparable to or modestly more vascularized than (-) (**Fig. 8**). A quantitative analysis of wound-bed angiogenesis (e.g., total blood-vessel length per unit area) is currently being performed with a standardized underside-imaging protocol.

**Figure 8.**
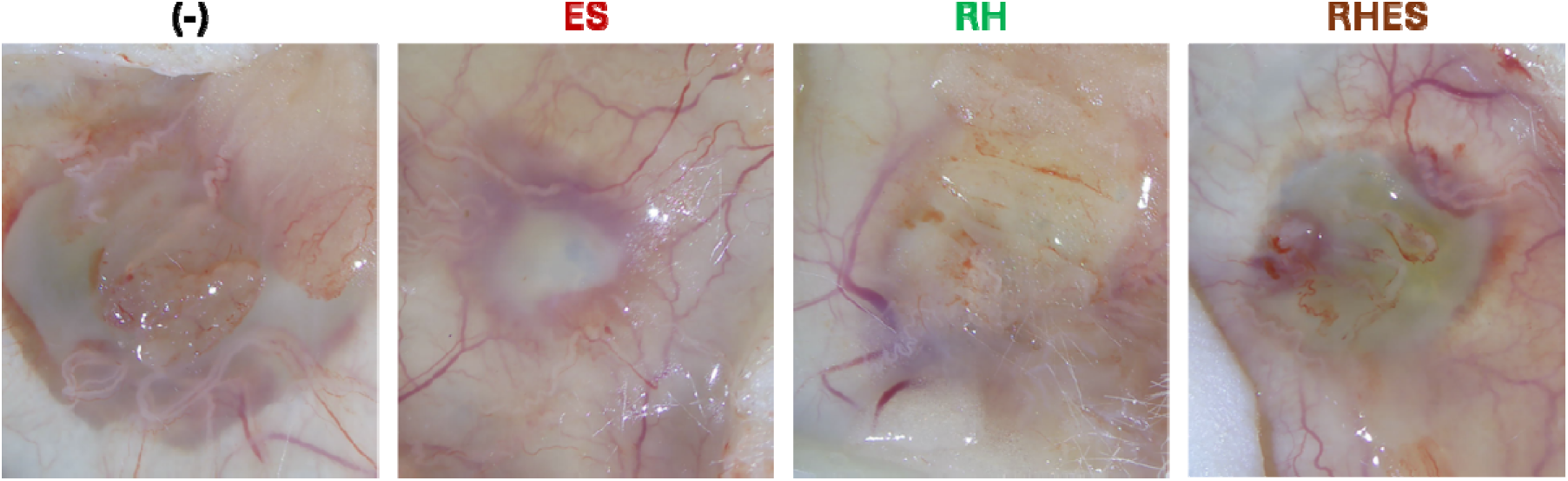
Representative underside of the wound images at day 7 showing the blood vessels involved in wound healing.

### 3.7 Regenerated tissue collagen content and cutaneous nerves

Masson’s trichrome (MT) staining showed the tricolor skin tissue cross-section at the interface of the intact skin and the regenerating wound site (**Fig. 9**). It is evident that those containing the residual hair treatment (RH and RHES) contain new connective tissues at the wound bed site. Collagen content in dermal granulation tissue measured by MT blue area fraction differed significantly across treatments (Kruskal-Wallis H = 16.2, **p = 0.001). RH and RHES wounds contained 6.1× and 8.5× more collagen, respectively, than (-) controls (median Mann-Whitney **p = 0.002 for both; Bonferroni-adjusted ***p = 0.007 for both). RH and RHES were both significantly higher than ES (***p = 0.009 for both pairwise). ES alone did not significantly differ from (-) in collagen content (p = 0.25) (**Fig. 10**).

**Figure 9.**
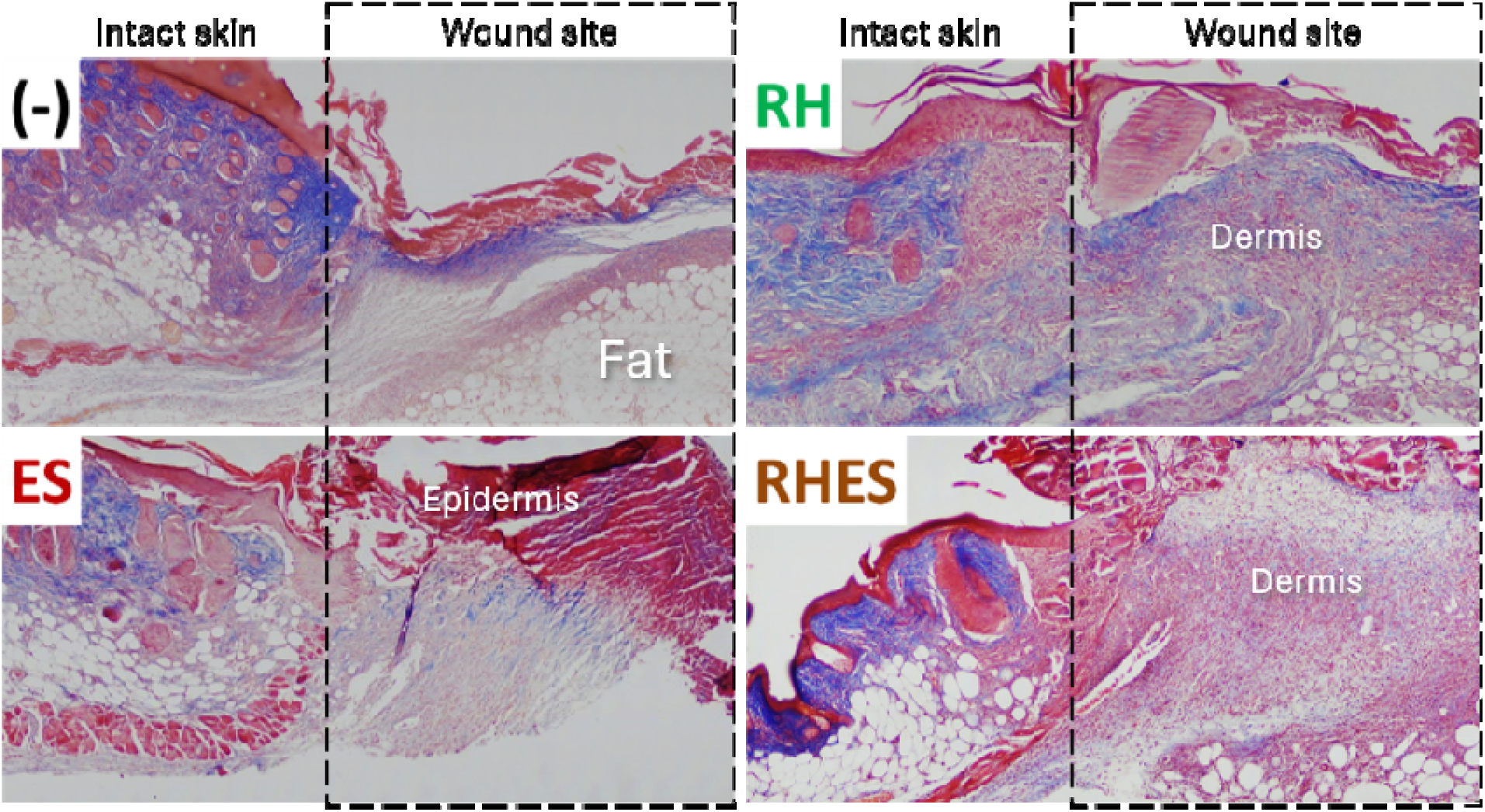
Representative Masson’s trichrome-stained cross sectional images showing the intact skin (left) and the wound site (right) per treatment group.

**Figure 10.**
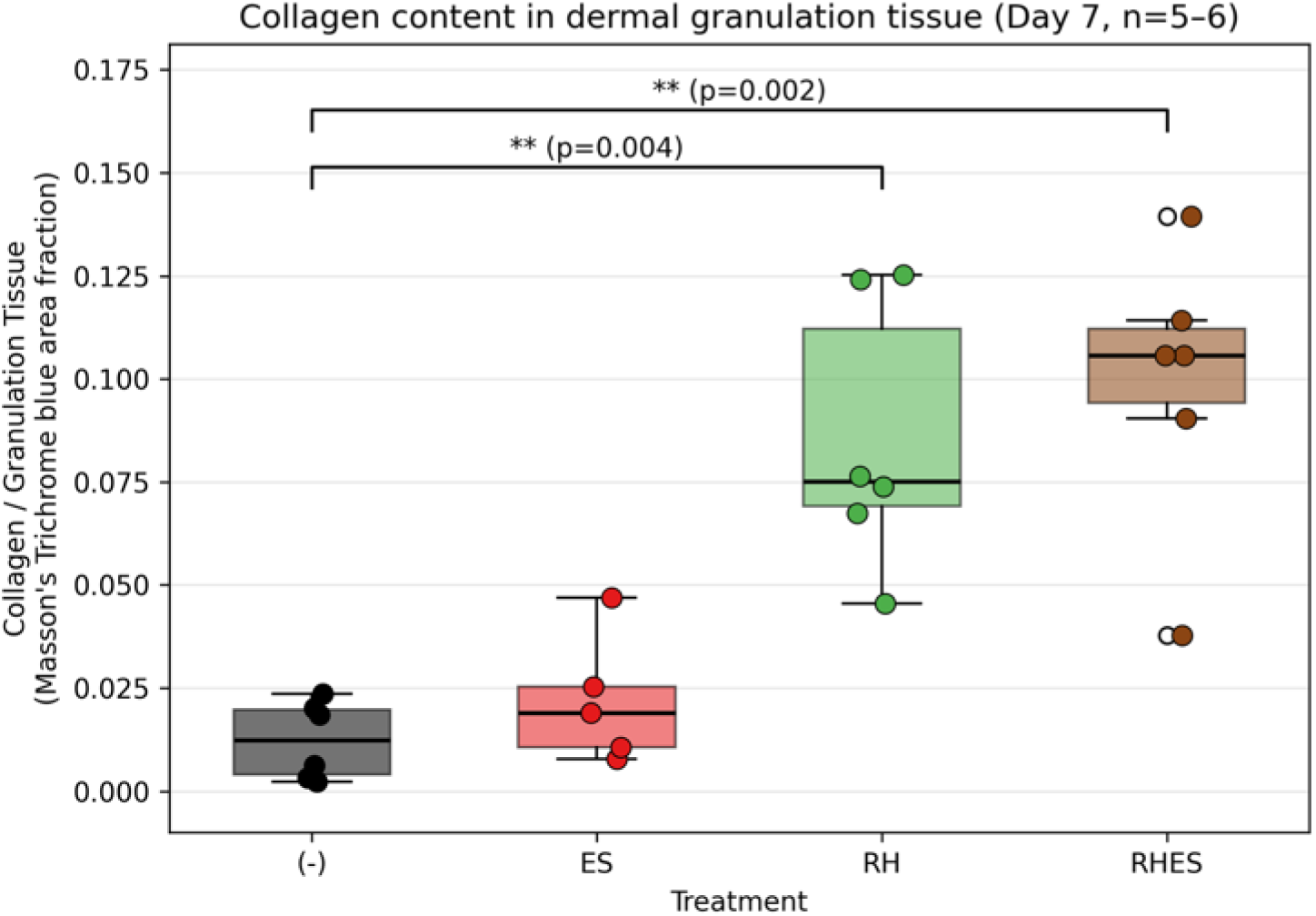
Collagen content in dermal granulation tissue at day 7 (MT blue area fraction within manually-traced granulation tissue ROI). Box: median and IQR; whiskers: 1.5 IQR; dots: individual wound sections. n per group: (-) n = 6, ES n = 5, RH n = 6, and RHES n = 6.

IHC staining displayed relatively higher GAP-43 signals for treatments with residual hair biomaterials (RH and RHES) (**Fig. 11**). GAP-43 expression in dermal granulation tissue measured by relative fluorescence intensity per tissue area also differed significantly across treatments (Kruskal-Wallis H = 15.6, **p = 0.001). RH and RHES wounds showed 16× and 27× greater mean GAP-43 expression, respectively, than (-) controls (Mann-Whitney *p = 0.020 and *0.014 respectively; Bonferroni-adjusted p = 0.058 and *0.042). RH and RHES were both significantly higher than ES (*p = 0.017 and **0.002 respectively). ES alone did not significantly differ from (-) in GAP-43 expression (p = 0.16) (**Fig. 12**).

**Figure 11.**
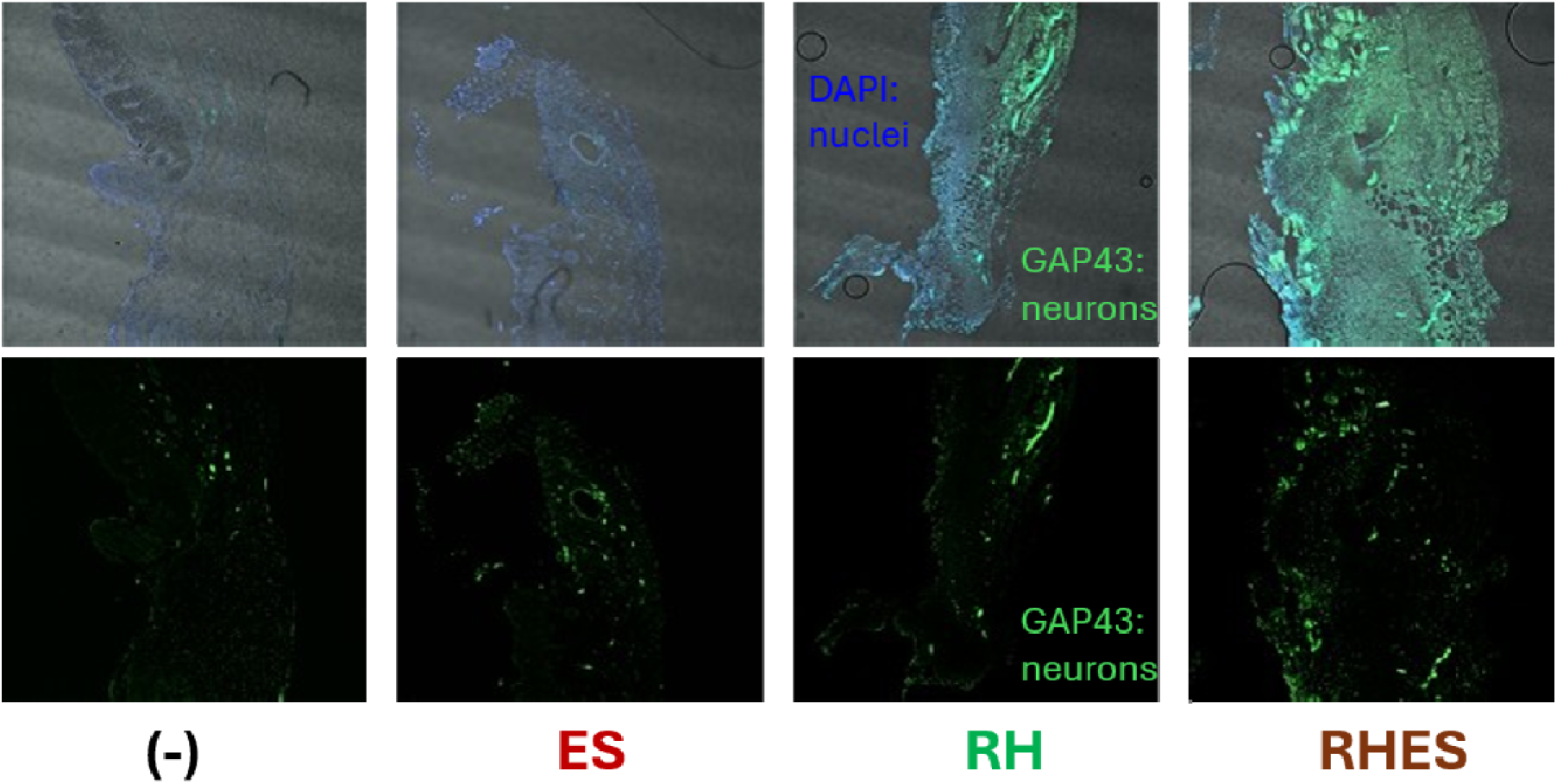
Representative GAP-43 immunostaining outcome showing DAPI-stained nuclei and overexposed images (top row) and GAP-43 only (bottom row) per treatment group.

**Figure 12.**
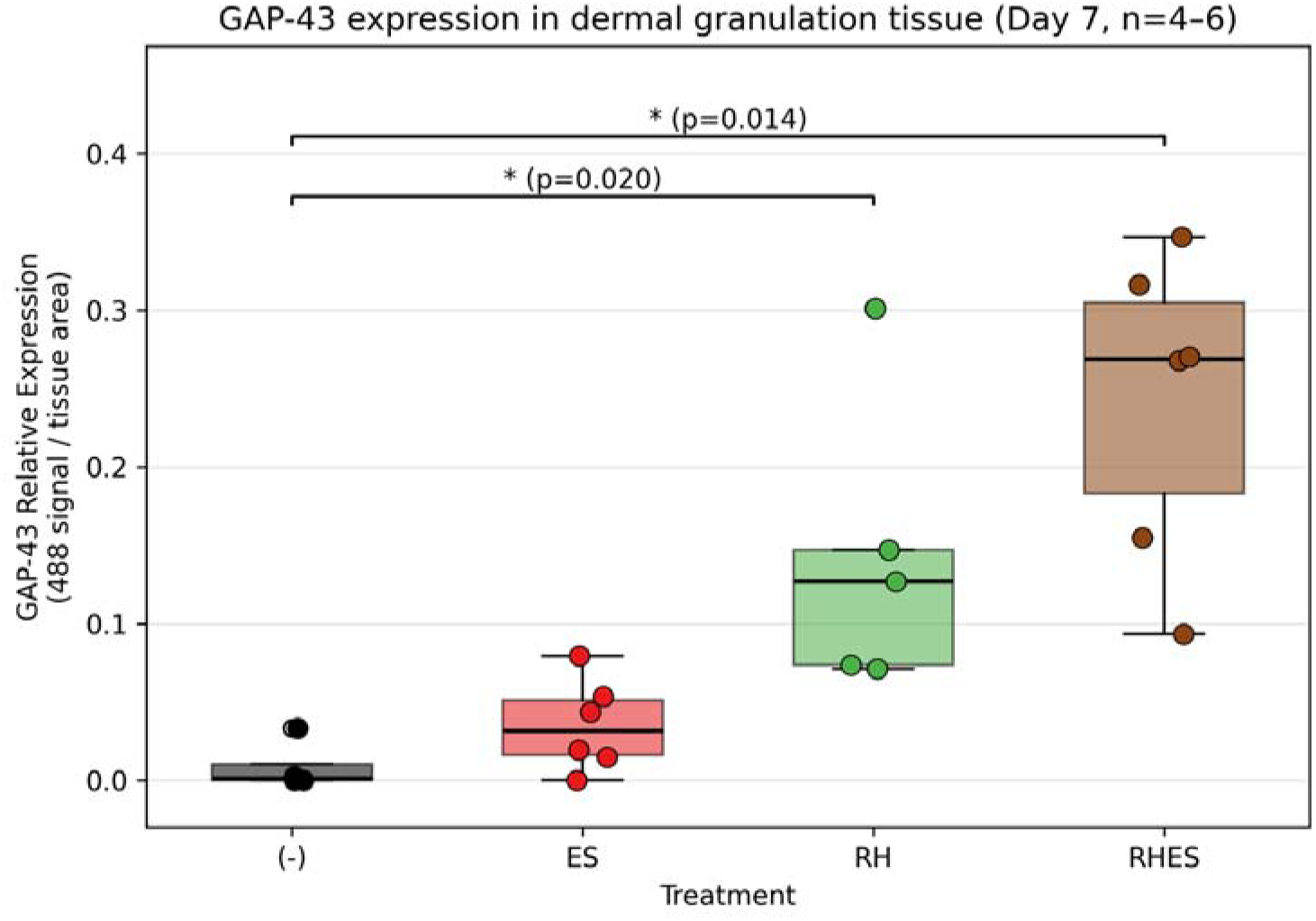
GAP-43 relative expression in dermal granulation tissue at day 7, measured as 488-nm-channel signal (green) per tissue area within a granulation tissue ROI. Box: median and IQR; whiskers: 1.5 IQR; dots: individual wound sections. n per group: (-) n = 4, ES n = 6, RH n = 5, and RHES n = 6. GAP-43 is a marker of regenerative cellular activity but is not unambiguously neuron-specific.

The MT and IHC GAP-43 results are concordant: both tissue markers identify RH and RHES as substantially elevated above (-) and ES, and both fail to distinguish ES from (-). This pattern, contrasting with the closure-kinetics finding in which ES showed the largest effect, suggests that ES and RH operate through distinct biological mechanisms during the proliferative phase of healing.

## 4. DISCUSSION

### 4.1 Effect of residual hair biomaterials, without or with electrical stimulation

This study evaluated wound size kinetics, wound shape, and tissue-level healing markers across four treatment conditions in a bilateral splinted murine excisional model. Three principal findings emerged: ES, RH, and RHES each significantly accelerate wound closure compared to untreated controls. RH and RHES significantly enhance collagen deposition and GAP-43 expression in dermal granulation tissue while ES alone does not. Wound shape is more circular and more stable in RH-containing groups. Together, these findings support that ES, RH, and RHES improve wound healing through partially distinct mechanisms, and that RH is the principal driver of tissue-level regenerative markers measured here.

### 4.2 Mechanistic interpretation

The contrast between the closure-kinetics ordering (ES > RHES > RH > (-)) and the tissue-marker ordering (RHES ≈ RH > ES ≈ (-)) is a central finding of this study. Two non-mutually-exclusive interpretations are consistent with the data. First, ES may act primarily through accelerated wound contraction and re-epithelialization, mechanisms that influence external wound dimensions but do not necessarily increase tissue regenerative activity over a 7-day window. Bioelectric fields are known to direct keratinocyte migration,^10,16,17^ and to modulate inflammation^11^, both of which would accelerate measured closure without directly elevating collagen synthesis or GAP-43 expression. Second, the RH biomaterial layer, by maintaining a moist wound environment and providing an organized keratin substrate^2,3^, may support fibroblast collagen synthesis and the cellular activity reflected by GAP-43 staining, independent of and additive to the closure-rate effects of ES. The combined RHES condition gains both benefits.

GAP-43 is associated with regenerating peripheral nerves and is a widely used marker of neuronal regrowth^18^, but it is also expressed by some non-neuronal cells in healing tissue. Without co-staining for a neuronal-lineage-specific marker (β-III tubulin or PGP9.5), we frame the present finding as elevated GAP-43 expression in dermal granulation tissue, a marker of regenerative cellular activity, rather than as a direct measurement of peripheral nerve regrowth. Co-stained follow-up work is warranted.

### 4.3 Significance for wound care

The mechanistic distinction is clinically relevant. Wound-care strategies that accelerate closure without correspondingly improving tissue quality may yield faster-closing wounds with weaker scars or impaired barrier function. The present data indicate that RH-containing treatments enhance two markers of tissue quality (collagen deposition and GAP-43 expression) that ES alone does not, while also improving wound shape stability. For wound types in which both speed and tissue quality matter, the combined RHES condition is the most attractive of those tested here, and a single-modality RH application may be preferred where speed is less critical than long-term tissue integrity.

### 4.4 Methodological contributions

Two methodological features of this study merit comment. First, the use of an AI-assisted review-and-correct workflow for boundary segmentation allowed a single expert tracer to validate boundaries on 288 images with substantial time saving (84.4% of predictions accepted without modification) while preserving full human review of every measurement entering the analysis. The workflow design, predictions reviewed in 8-day per-wound batches, with explicit ACCEPT-CORRECT-REJECT classification, is generalizable to similar image-analysis problems where ground-truth annotation is expensive. Second, the linear mixed-effects framework with per-image observation covariates allowed us to quantify and partially control for known sources of measurement artifact (wound dryness and eschar coverage) without losing power from list-wise exclusion of affected images. Two covariates, Dryness and EscharCoverage, emerged as significant negative predictors of measured wound size, and the treatment effects survive their inclusion.

### 4.5 Limitations

The histological subsample (6 mice per group) is small, so individual histological group means have wide confidence intervals. We report mean-based fold-changes and non-parametric tests to reflect this. For GAP-43, the near-zero control values make fold-change estimates sensitive to the choice of central tendency (the corresponding median-based ratios are substantially larger, ∼ 100× and ∼ 200×), so the significance tests rather than the fold magnitudes should be regarded as the primary evidence. Three histology sections (one each in ES MT, plus two in (-) GAP-43 and one in RH GAP-43) were excluded due to processing damage or unreadable staining. This missingness reduces the n further in those samples. The 7-day endpoint captures the early proliferative phase but does not extend to complete re-epithelialization or scar maturation, so the long-term consequences of the closure-kinetics and tissue-quality differences observed here remain to be characterized. The wound shape findings are descriptive. No formal mechanical testing was performed. Finally, this study used a single expert tracer for the AI training set and for review of automated predictions. Multi-rater agreement on the AI training labels would further strengthen the methodology.

## 5. CONCLUSIONS

In a bilateral splinted murine excisional wound model with an excipient-tuned RH formulation and verified ES current, all three treatments significantly accelerated wound closure compared to untreated controls (Day × Treatment ***p < 0.0001), with ES showing the largest closure-rate advantage. RH and RHES additionally enhanced collagen deposition (6× and 8×, respectively; both significant after Bonferroni correction) and GAP-43 expression (16× and 27× greater mean expression, respectively); the GAP-43 increase remained significant after correction for RHES (adjusted p = 0.042) but not for RH (adjusted p = 0.058). ES alone did not significantly increase either tissue marker. These findings support RH biomaterial as a wound-care strategy that improves tissue-level outcomes during the early proliferative phase, and indicate that the combination with ES achieves both faster closure and improved tissue regeneration.

## AUTHOR CONTRIBUTIONS

Dana Saparova: conceptualization, methodology, formal analysis, investigation, data curation, writing - original draft, review & editing, visualization. Zara Mahmood: investigation. Hope Samuel: investigation. Jessica Barayuga: investigation. Jash Mody: investigation. Nancy Radecker: investigation. Roche C. de Guzman: conceptualization, methodology, software, formal analysis, investigation, resources, data curation, writing - review & editing, visualization.

## ACKNOWLEDGEMENTS

We thank Esther Thomas for assistance with murine surgery and monitoring; Dr. Chris Boyko and the Hofstra Biology Animal Facility for animal care; the Bioengineering Materials Lab of Dr. de Guzman; the Rabinowitz Honors College for research assistant funding; and the Hofstra Business School Innovation Award for partial materials, supplies and support.

## ETHICAL STATEMENT

All animal procedures were approved by the Hofstra University Institutional Animal Care and Use Committee and conducted in accordance with NIH and institutional guidelines for the care and use of laboratory animals.

## DATA AVAILABILITY

De-identified raw image data, ImageJ ROI files, manual measurements, and analysis code are available from the corresponding author on reasonable request. Subject to patent counsel review prior to public deposition.

